# Bilateral Symmetry and Asymmetry in the C. elegans Connectome: A Graph-Theoretic Analysis based on Redundancy Measures

**DOI:** 10.1101/2024.10.03.616419

**Authors:** Pyeong Soo Kim, Youngjo Song, Jerald D. Kralik, Jaeseung Jeong

## Abstract

Understanding the balance between symmetry and asymmetry in animal nervous systems is crucial for unraveling the complexities of neural architectures and their functions. Previous studies have primarily focused on morphological symmetry, such as neuron placement, leaving the symmetry in the functional architecture largely unexplored. The current study investigates this aspect within the *Caenorhabditis elegans* connectomes by introducing a graph-theoretic approach. By defining a ‘mirror-symmetry index,’ we quantitatively assess the symmetry in these connectomes, revealing a significant level of bilateral symmetry alongside notable asymmetry. Our approach also incorporates measures including connectivity similarity, motif-fingerprint differences, and path-compensation index to evaluate the network’s functional redundancy and its capacity to compensate for unilateral disturbances. Here we show the C. elegans connectomes’ robust bilateral symmetry, which not only facilitates similar functions across neuron pairs but also ensures resilience against disruptions. This redundancy is not confined to symmetrical connections; it also includes asymmetric ones, adding to the neural network’s complexity. An in-depth analysis into different neuron types shows varied redundancy levels: high in interneurons, moderate in motor neurons, and low in sensory neurons. This pattern suggests a strategic neural design where diverse inputs from sensory neurons, coupled with the stable integration by interneurons, lead to coordinated actions through motor neurons. This study advances our understanding of neural connectomes, offering insights into the intricate balance of symmetry and asymmetry in neural systems and their implications for complex, adaptive behaviors.

## Introduction

*Bilateral symmetry* in the nervous system, displaying nearly identical mirror images when divided along the sagittal plane, is a characteristic observed widely in the group of animals known as *Bilateria*. This group emerged about 555 million years ago (Evans et al., 2020) and includes a variety of animals often studied in neuroscience (about 99% of all animals) (Finnerty et al., 2004), such as humans, mammals, insects, and nematodes like *Caenorhabditis elegans* (*C. elegans)*. The prevalent nature of such symmetry signifies the potential operational benefits from such structural configurations. Indeed, many theoretical and empirical studies have shown that network symmetry (not exclusive to bilateral symmetry) results in redundancy (i.e., the replication of identical components), thereby influencing various network dynamics, such as robustness (Dekker and Colbert, 2004, Tononi et al., 1999, Albert and Barabási, 2002), controllability (Chapman and Mesbahi, 2015, Whalen et al., 2015, Yuan et al., 2013), and synchronization (Nicosia et al., 2013, Nishikawa and Motter, 2016, Avila et al., 2023, Pecora et al., 2014). In particular, robustness — a system’s ability to sustain stability and reliability in the face of disruptions (Demetrius and Manke, 2005, Kitano, 2004) — is an essential attribute of any complex evolving system (Kitano, 2004). This ability can be strengthened by symmetry since symmetry induces redundancy, which offers backup mechanisms or alternative routes using identical or similar elements, ensuring the continuity of essential functions even if specific system components are disrupted or fail (Kitano, 2004), thus suggesting the evolutionary benefit of having a large-scale symmetry, such as the bilateral symmetry.

Despite the benefits of bilateral symmetry, the Bilaterians’ nervous system also accommodates asymmetry (Hugdahl, 2005, Toga and Thompson, 2003), leading to lateralization of brain functions (i.e., functional asymmetry): e.g., left-hemispheric dominance of humans’ language processing (Geschwind, 1970, Vigneau et al., 2006), the right hemisphere’s advantage in face recognition tasks in chimpanzees and sheep (Morris and Hopkins, 1993, Peirce et al., 2000), and left-hemisphere advantage in birds’ stimuli categorization (Rogers, 2014, Vallortigara, 1989, Xiao and Güntürkün, 2018). In fact, perfect (bilateral) symmetry, leading to complete redundancy, is not ideal in real biological contexts since large-scale replication of neural elements can overtax metabolic resources to perform a single function, thus compromising diversity and flexibility (Kitano, 2004). Frequent observations show that departure from perfect bilateral symmetry (i.e., brain lateralization) is associated with cognitive flexibility or capacity (Rogers, 2021). For instance, lateralization in domestic chicks has been linked with advanced multitasking capabilities (Rogers et al., 2004, Wichman et al., 2009); even invertebrates with significant side-biases, suggestive of brain lateralization, exhibit enhanced learning abilities and fewer reaching errors (Miler et al., 2017, Bell and Niven, 2016). However, lateralization comes with a trade-off: a lateralized brain may be less robust in stressful stimuli (Byrnes et al., 2016, Barnard et al., 2016).

In sum, the symmetry and asymmetry in nervous systems might reflect a trade-off or balance between redundancy and functional diversity (for review, see Rogers, 2021). Referred to as *degeneracy*, this balance might be the most viable solution for evolving biological systems, as repeating identical elements can be costly and can constrain functional diversity and complexity (Kitano, 2004, Tononi et al., 1999, Noppeney et al., 2004). Therefore, understanding the balancing of these two counterpoint designs could illuminate the fundamental design objectives of the nervous system. However, the intricate details and functional implications of symmetry and asymmetry in the nervous system at the neuronal level remain largely unexplored, primarily due to the complexity of the interaction between these two phenomena (i.e., redundancy and functional diversity). For this reason, most research on network symmetry has focused on studying perfect symmetries, encapsulated by the concept of *automorphism* in algebraic graph theory (MacArthur et al., 2008, Chapman and Mesbahi, 2015, Nicosia et al., 2013, Whalen et al., 2015, Nishikawa and Motter, 2016, Avila et al., 2023, Pecora et al., 2014, Sánchez-García, 2020); consequently, it is challenging to directly apply those works to the exploration of bilateral symmetry in the nervous system, which is inherently imperfect. Nonetheless, recent studies addressed this issue from various perspectives. For instance, Morone and Makse (2019) proposed the idea of *pseudosymmetry*, which offers a more flexible take on the concept of automorphism. They used this measure as an alternative to perfect symmetries (i.e., automorphism) in the connectome, aiming to identify robust functional building blocks even with minor changes in connections. Although pseudosymmetry presents a promising method to study large-scale symmetry (such as bilateral symmetry) by acknowledging its inherent imperfections (such as asymmetry), the scope of their research was limited to specific subnetworks like forward and backward motor networks. Importantly, bilateral (a)symmetry is yet to be comprehensively explored. Meanwhile, another study by Pedigo et al. (2023) investigated bilateral asymmetry by introducing a statistical framework for its evaluation. They demonstrated significant global differences (i.e., connection densities) between the bilateral aspects of the *D. melanogaster* larva’s connectome, thus providing valuable insight into how bilateral asymmetry can be statistically assessed. However, their approach largely ignored the local topological properties (e.g., wiring patterns) due to the primary focus on the global aspect, thereby leaving the connectome functionality for future study.

Therefore, the current study aims to investigate the intricate nature of bilateral (a)symmetry within the connectome’s functionalities and to assess the influence of the bilaterally symmetric structure on the functionalities using various graph-theoretic network measures. For this investigation, we use the *C. elegans* connectome, as its completely mapped and straightforward nervous system offers a unique opportunity to probe the existence and interplay of symmetry and asymmetry. Fundamentally, the *C. elegans* nervous system displays substantial symmetry: a large portion of the neurons are arranged into pairs of bilaterally symmetrical neurons (see Figure 1A), with detailed symmetric aspects like symmetric synaptic connectivity; furthermore, the bilaterally paired neurons often share similar developmental paths, indicating lineage-based symmetry (Sulston et al., 1983, White et al., 1986, Hobert et al., 2002). Yet, despite the pronounced symmetry, the *C. elegans* connectome also exhibits significant asymmetry, which is evidenced by a significant ratio of bilateral neurons’ distinct left and right connections (Cook et al., 2019) and the presence of two functionally lateralized, bilateral neuron pairs: ASE and AWC neurons (Hobert et al., 2002, Downes et al., 2012, Hobert, 2014). *ASE neurons*, the main taste receptor of *C. elegans*, show asymmetric roles; the left ASE neuron detects Na+, and the right one senses Cl- and K+, due to different gene activations (Ortiz et al., 2009, Harrell and Goldstein, 2011). On the other hand, AWC neurons function in odor detection, with only one randomly activated to detect specific odors, an event regulated by gene expression (Bauer Huang et al., 2007). Given these findings, we leveraged the extensive structural and functional data to develop and apply graph-theoretic measures for probing the bilateral symmetry and asymmetry within the nervous system.

**Figure 1.**
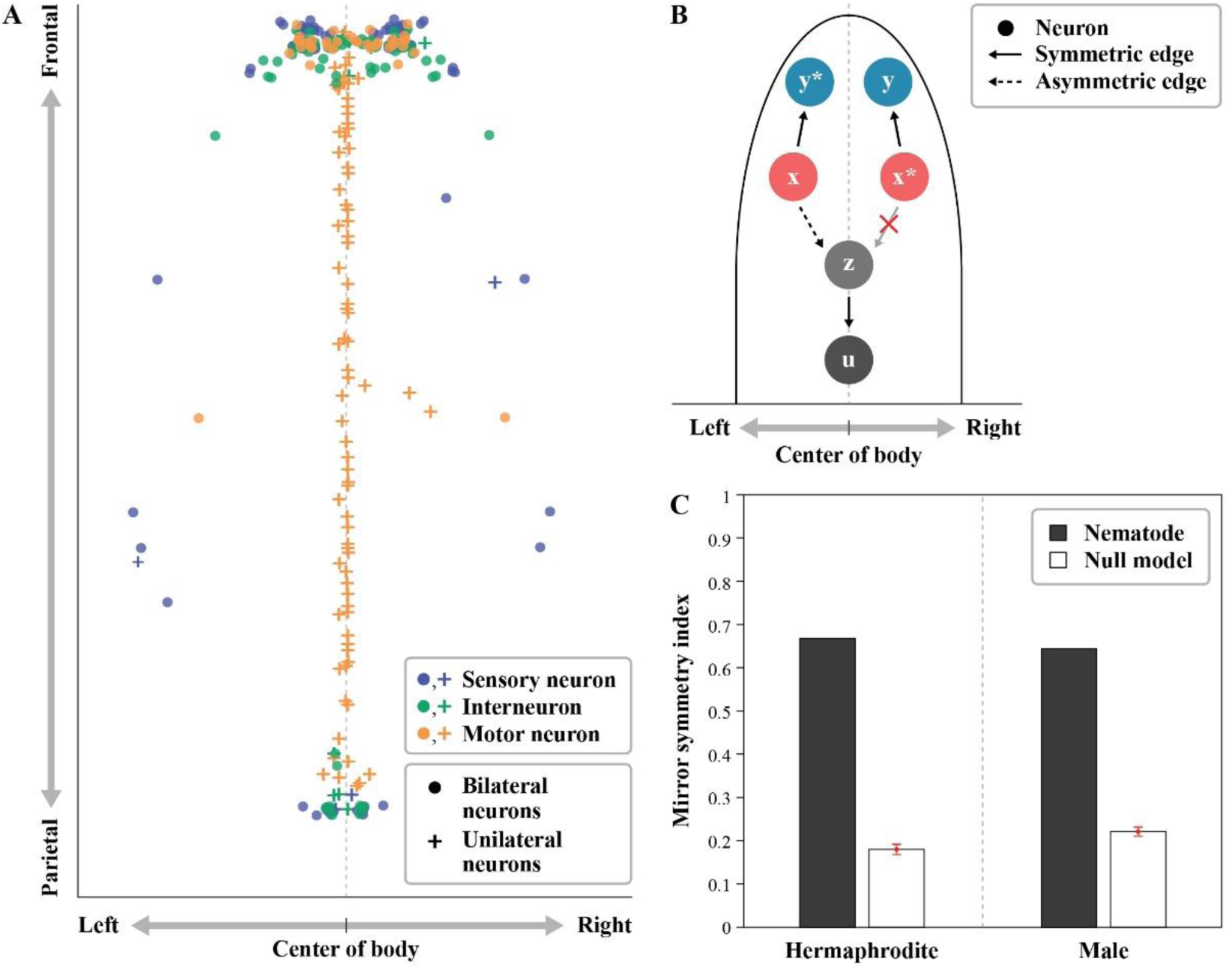
**(A)** The spatial configuration of the hermaphrodite *C. elegans* neuron’s soma. In a bilaterally symmetric pair, the left neurons are denoted by the letter ‘L’ and the right neurons by ‘R’. Unilateral neurons are denoted by a plus sign ‘+’. The sensory neurons are shown in blue, interneurons in green, and motor neurons in orange. Note that around 69% of motor neurons are unilateral, mostly located along the center of the body (i.e., sagittal plane). (B) **Defining contralateral neurons, edges, and mirror-symmetric edges.** Contralateral neurons are denoted by an asterisk (*). The edges, (*x*, *y*\*), (*x*\*, *y*), and (*z*, *u*), have contralateral connections (*x*\*, *y*), (*x*, *y*\*), and (*z*, *u*), respectively. Meanwhile, the edge (*x*, *z*) have no contralateral connection. (C) **The mirror-symmetric indices for both hermaphrodite and male *C. elegans***. These values were compared against null models (i.e., random networks). A total of 1,000 null models were created, preserving the degree sequence in the process (see Methods).

More specifically, the current investigation was conducted via three main steps. (1) In the first step (Results section 1), we introduced the “mirror-symmetry index,” which allows us to measure the level of bilateral symmetry within the *C. elegans* connectome networks while also recognizing and accounting for any asymmetry that may exist. (2) In the second phase (Results sections 2-4), we applied three distinct graph-theoretic measures—comprising one established and two novel metrics—to capture various dimensions of functional redundancy: (a) *connectivity similarity*, illustrating redundancy as a similarity in the input-output profiles (which has been used to reflect similarity of neuronal functions in previous studies), (b) *motif-fingerprint difference*, reflecting redundancy as similarity in local operations between bilateral sides, and (c) *path-compensation index*, which quantifies the ability of one side to compensate for the impairment of its counterpart by providing an alternative pathway for global information processes. This multifaceted assessment was necessary because, unlike the previous studies using automorphism, biological systems rarely exhibit perfect bilateral symmetry, complicating the clear identification of identical components between bilateral sides. Consequently, the concept of redundancy becomes dependent on the level of abstraction (i.e., what specific details are ignored) employed in the evaluation, with each level reflecting a particular aspect of the network’s functionality. (3) The final step (Results section 5) focused on investigating the bilateral redundancy within subsystems, such as sensory, motor, and interneuron, and also exploring the redundancy between each pair of bilateral neurons, allowing for an in-depth understanding of the design objectives within the network.

## Results

In the current study, we analyzed the connectomes of both hermaphrodite and male *C. elegans*, which were created by Cook et al. (2019). These connectomes provide detailed information on chemical synapses and gap junctions, including their direction, type, and strength (Sohn et al., 2011). By examining connectomes from both sexes, we aimed to gain a thorough understanding of neural structures and functions at the cellular level. This approach allowed us to identify unique and shared traits, compare neural organization, and validate findings across different neural and behavioral contexts.

Our method involved representing the connectome as a directed binary graph, concentrating on whether connections exist or not. This simplification has proven effective in uncovering basic pathways and structures in the brain, while still retaining the critical characteristics of the network dynamics (Varshney et al., 2011, Towlson et al., 2013, Watts and Strogatz, 1998, Uzel et al., 2022). Utilizing the provided connectome data, we generated a directed binary graph (G) for each sex of *C. elegans* - one for the hermaphrodite and one for the male. We assigned each neuron as a vertex (or node) in the graph, indicated by lower alphabets (e.g., *x*, *y*). The vertex set of the graph G is denoted as V. Information about connections was used to draw edges between these nodes, considering the direction of synapses and gap junctions while disregarding details such as the types or strengths of connections. This meant that if neuron *x* connected to neuron *y*, an edge was drawn from *x* to *y*, notated as (*x*, *y*). If there was no connection, no edge was drawn. The edge set of the graph G is denoted as E.

Furthermore, drawing upon the nomenclature introduced for *C. elegans* (White et al., 1986), we applied descriptive labels to the nodes within the graph, designating them as either *bilateral neurons* or *unilateral neurons*. Bilateral neurons exist in pairs - one on the left and one on the right, and each neuron is noted by the suffix L/R. For example, ASEL represents the left ASE neuron, while ASER represents the right ASE neuron. Neurons that do not form pairs and exist only on one side or close to the sagittal plane are called unilateral neurons. Most unilateral neurons were located along the sagittal plane, with some exceptions (refer to Figure 1A).

### Assessing a high degree of mirror symmetry in *the* C. elegans *connectome*

To establish the concept of bilateral symmetry, we initially draw upon the conventional understanding of *mirror symmetry* - where the left and right sides of a system are exact reflections of each other (Hugdahl, 2005, Hobert et al., 2002, Watkins et al., 2001, Takao et al., 2011, Good et al., 2001). However, in the case of the *C. elegans* connectome, and indeed any brain network, it is important to recognize that the left and right sides are not perfect mirror images due to the presence of bilateral asymmetry. Consequently, our notion of mirror symmetry has to be relaxed to accommodate these asymmetries. Instead of requiring perfect reflection, we aim to quantify the degree of bilateral symmetry, i.e., how much the network displays bilateral symmetry. In other words, we are interested in measuring how closely the left and right sides of the network resemble each other, acknowledging that they may not be identical.

With this in mind, we introduce the concept of a “*contralateral neuron*,” which refers to a neuron’s counterpart on the other side (left or right) of the network (Figure 1B). We designate these contralateral neurons with an asterisk (*). For instance, in a pair of bilateral neurons like the ASE neurons, if we take *x*=ASEL (ASE left), its contralateral neuron would be *x*\*=ASER (ASE right). Similarly, if we consider *y*=AWCR (AWC right), its contralateral neuron is *y*\*=AWCL (AWC left). For unilateral neurons, which do not have a counterpart on the other side, we define their contralateral neuron as themselves. For example, for a unilateral neuron, like *z*=PVT, its contralateral neuron is defined as *z*\*=*z*=PVT.

More significantly, we extend this idea to “contralateral edges,” which refers to counterpart connections or edges on the opposite side of the network, also denoted with an asterisk (*). Essentially, a contralateral edge of a given edge can be characterized as a directional connection between the contralateral neurons of the pair connected by the given edge. For example, if we have an edge (*x*, *y*)=(ASEL, AWCR) linking ASEL and AWCR, its contralateral edge would be (*x*, *y*)*=(*x*\*, *y*\*)=(ASER, AWCL), connecting ASER and AWCL. Furthermore, we classify an edge as “mirror symmetric” if its contralateral edge exists in the network, i.e., if (*x*, *y*)* is an element of the edge set *E*(*G*) ((*x*, *y*) ∈ *E*(*G*)). If the contralateral edge does not exist in the network, we classify the edge as “mirror asymmetric.”

Building on these classifications, we propose the “mirror-symmetry index,” calculated as the ratio of symmetric edges to the total number of edges in the network. If the network is entirely symmetric (meaning the left side mirrors the right side perfectly), the mirror-symmetry index equals one. Conversely, if the network is entirely asymmetric (having no contralateral edges), the mirror-symmetry index is zero. Therefore, a higher mirror-symmetry index implies a greater extent of mirror symmetry in the network. Note that the mirror-symmetry index can be understood as a measure of the extent of left-right automorphism (i.e., invariance under the left-right label permutations of all vertices), which also can be seen as pseudosymmetry (Morone and Makse, 2019) regarding the left-right label permutations of all vertices.

We examined the *C. elegans* connectome networks and found a pronounced presence of mirror symmetry in both sexes. For hermaphrodite and male connectomes, the mirror-symmetry indices were determined to be 0.68 and 0.66, respectively. These mirror-symmetry indices were markedly larger than those achievable in comparable random networks (retaining the same degree distributions, n = 1,000; refer to Methods). The corresponding mean mirror-symmetry indices for these random networks were only 0.18 (with a standard deviation of 0.0060) for networks rewired using the hermaphrodite connectome and 0.23 (with a standard deviation of 0.0051) for those rewired using the male connectome (as shown in Figure 1C). This significant discrepancy indicates the pronounced structural similarity between the left and right sides of the *C. elegans* connectomes, implying a likely existence of bilateral redundancy in their network functions.

It is, however, crucial to point out that despite the high levels of mirror symmetry, the mirror-symmetry indices were considerably smaller than one (i.e., ∼0.7); thus, around 30% of the connections exhibited mirror asymmetry. This substantial asymmetry suggests that the two sides of the network are not completely redundant.

### Both mirror-symmetric and mirror-asymmetric edges enhance connectivity similarity

In line with our expectations, the *C. elegans* connectomes demonstrated a high level of mirror symmetry. This pronounced symmetry may lead to redundancy across the bilateral sides, potentially enabling them to display identical or at least comparable functions. One commonly used measure of (functional) redundancy is *connectivity similarity,* which measures how much the connection profiles of two neurons overlap, as illustrated in Figure 2A. This measure is sometimes referred to by other names, such as the matching index, Jaccard index, or connectivity fingerprints, as per various studies (Passingham et al., 2002, Hilgetag et al., 2002, Zamora-López et al., 2010, de Reus and van den Heuvel, 2013, Fornito et al., 2016). This measure has proven useful in assessing the structural equivalence of pairs of neurons (Fornito et al., 2016); and Uzel et al. (2022) showed that the similarity in input connectivity, particularly 2-step similarity — how similar are the 2-step neighbors (Yip and Horvath, 2007) between two given neurons — correlates to a neuronal activation correlation between these neurons (with a correlation of about 0.5). In simpler terms, if two neurons have similar connections with other neurons, they are likely to have similar functions. Based on these findings, we opted to use connectivity similarity as one of our measures to evaluate bilateral redundancy in the *C. elegans* connectome (see Methods). Upon investigation, we found that the connectivity similarities, considering both 1-step and 2-step, as well as input similarity and input-output similarity, were significantly higher in bilateral neurons than in any possible pairs of neurons (all neurons) (p<0.0001 for all comparisons; see Figure 2B). Additionally, the connectivity similarities of the bilateral neurons were higher than those in the bilateral neurons of the null models (i.e., random networks) (p<0.001 for all comparisons; see Figure 2B). This finding indicates a noteworthy amount of bilateral redundancy in the network functions (Uzel et al., 2022).

**Figure 2.**
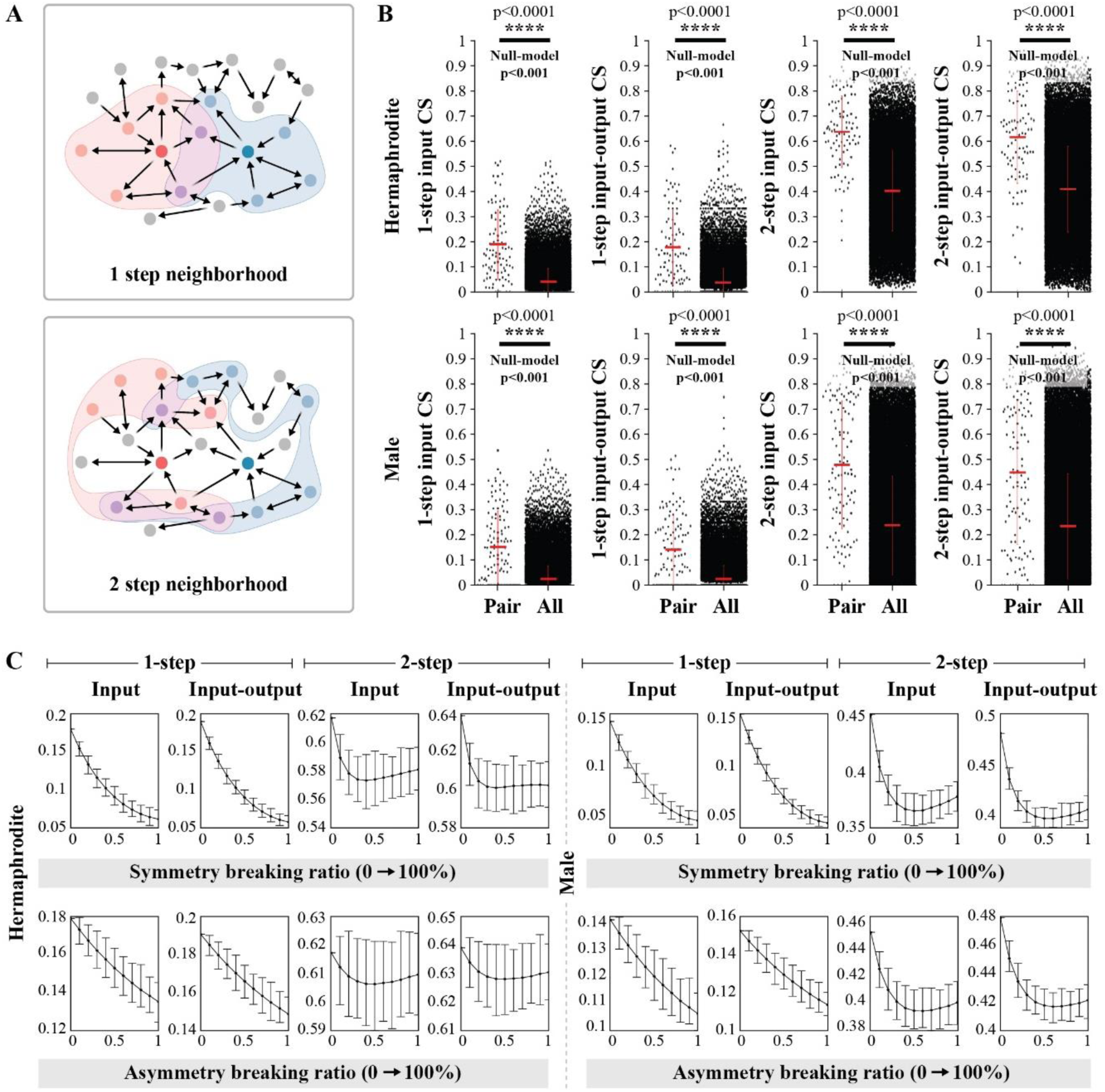
**(A)** An illustration of connectivity similarity. Connectivity similarity quantifies the overlap of neighborhoods in relation to the input-output connection. The determination of one-step or two-step similarity is based on the distance considered for the neighboring connections (compare one-step neighborhood and two-step neighborhood). (B) **Connectivity similarity of hermaphrodite and male *C. elegans*.** In all eight comparisons depicted in this figure, bilateral pairs show a higher connectivity similarity than all pairs (custom shuffle test, p<0.0001). The connectivity similarities are also significantly higher than random networks (null models, p<0.001). The red error bar indicates the mean and standard deviation. The null-model result compares the connectivity-similarity values between bilateral pairs in a randomly shuffled network, where the degree sequence is maintained, and bilateral pairs in the actual network of *C. elegans*. (C) **Connectivity** similarity results of symmetric and asymmetric edge shuffling for hermaphrodite and male *C. elegans*.

Following this observation, we further examined whether this significant similarity was exclusively the result of mirror-symmetric edges or not. To assess the influence of these from the overall connectivity similarity, we preserved all mirror-symmetric edges while we introduced variability in the mirror-asymmetric edges (i.e., connections unique to one side) through shuffling and vice versa (preserving asymmetric, varying symmetric). This process was performed in varying proportions to observe its effect on connectivity similarity (see Methods section for more detail).

Here, we found that with an increase in the shuffling ratio of the mirror-symmetric edges, while retaining all the mirror-asymmetric edges, the connectivity similarity consistently decreased (Figure 2C, yet there is an exception in the 2-step connectivity similarity of the hermaphrodite; see next paragraph), indicating that the high mirror symmetry enhances redundancy. Intriguingly, we also found that with an increase in the shuffling ratio of the mirror-asymmetric edges, while retaining all the mirror-symmetric edges, the connectivity similarity also decreased comparably (Figure 2C, yet there are again exceptions in the 2-step connectivity similarity of the hermaphrodite; see next paragraph). This finding was somewhat unexpected since it suggests that the asymmetric connections also enhance functional redundancy between bilateral neurons.

In the case of the hermaphrodite, however, we found that the decrease in 2-step connectivity similarity was not significant (in particular, 2-step input similarity in symmetry breaking and 2-step input similarity & input-output similarity in asymmetry breaking), as shown in Figure 2C. Yet, when all edges were shuffled (regardless of whether they were mirror-symmetric or not, i.e., null models), the 2-step connectivity similarity significantly decreased (see Figure 2B). This finding indicates that maintaining either type of edge (mirror-symmetric or asymmetric) is sufficient to preserve the 2-step connectivity, implying that both kinds of edges contribute to the redundancy between bilateral neurons. In sum, these findings underscore that making mirror-symmetric connections is not the sole design strategy determining the bilateral redundancy, suggesting the existence of other underlying principles contributing to enhancing bilateral redundancy.

### The motif fingerprints indicate degeneracy between bilateral neurons

A limitation of the connectivity similarity lies in its primary focus on the connection profiles (the identities of sources or destinations), which overlooks the distinct operations in which individual neurons may be involved. This means that two neurons might be engaged in different operations (such as synchronization, integration, or signal relay) even when they share the same signal inputs (due to overlapping connectivity). *Motifs*, which are local patterns of connections (see Figure 4A), can help address this limitation. Motifs act as foundational building blocks of networks, and their variations can reflect differing functional roles and network topology (Milo et al., 2004, Milo et al., 2002). Computational studies have shown that different types of motifs can influence specific functional roles, such as maintaining network activity and managing synchronization (Gollo and Breakspear, 2014, Gollo et al., 2014, Garcia et al., 2014), thus motifs have been employed as a tool for understanding the structure, functionality, and process of information integration in brain networks (Battiston et al., 2017, Morgan et al., 2018, Sporns and Kötter, 2004). Thus, building upon the premise that motif configuration (motif fingerprint) reflects the network functions, we aimed to deduce the functional difference between bilateral neurons (i.e., potential functional lateralization) that might not be inferred from connectivity similarities by analyzing the differences in their motif fingerprints.

Furthermore, motif analysis can provide insights into the crucial interplay of symmetric and asymmetric edges in functional lateralization. For instance, a 3-node motif can be any of 13 possible configurations of at least two connections among the three interconnected nodes (Figure 3A). Looking at the motifs for bilateral neuron pairs, depicted in Figure 3B, where each neuron is part of motifs *m*^3^_3_ and *m*^3^_6_, there are two mirror-symmetric connections within each motif, and an additional mirror-asymmetric connection present in *m*^3^. This arrangement has a high mirror-symmetry ratio of 0.8 since four out of five connections are mirror-symmetric. However, previous research has shown that these two motifs have different circuitry properties: *m*^3^_3_ exhibits largely synchronized activity, while *m*^3^_6_ does not (Gollo et al., 2014). Thus, a high symmetry level does not guarantee the functional similarity between bilateral sides of networks. This underlines the importance of motif analysis as a tool for investigating functional redundancy in the presence of asymmetry.

**Figure 3.**
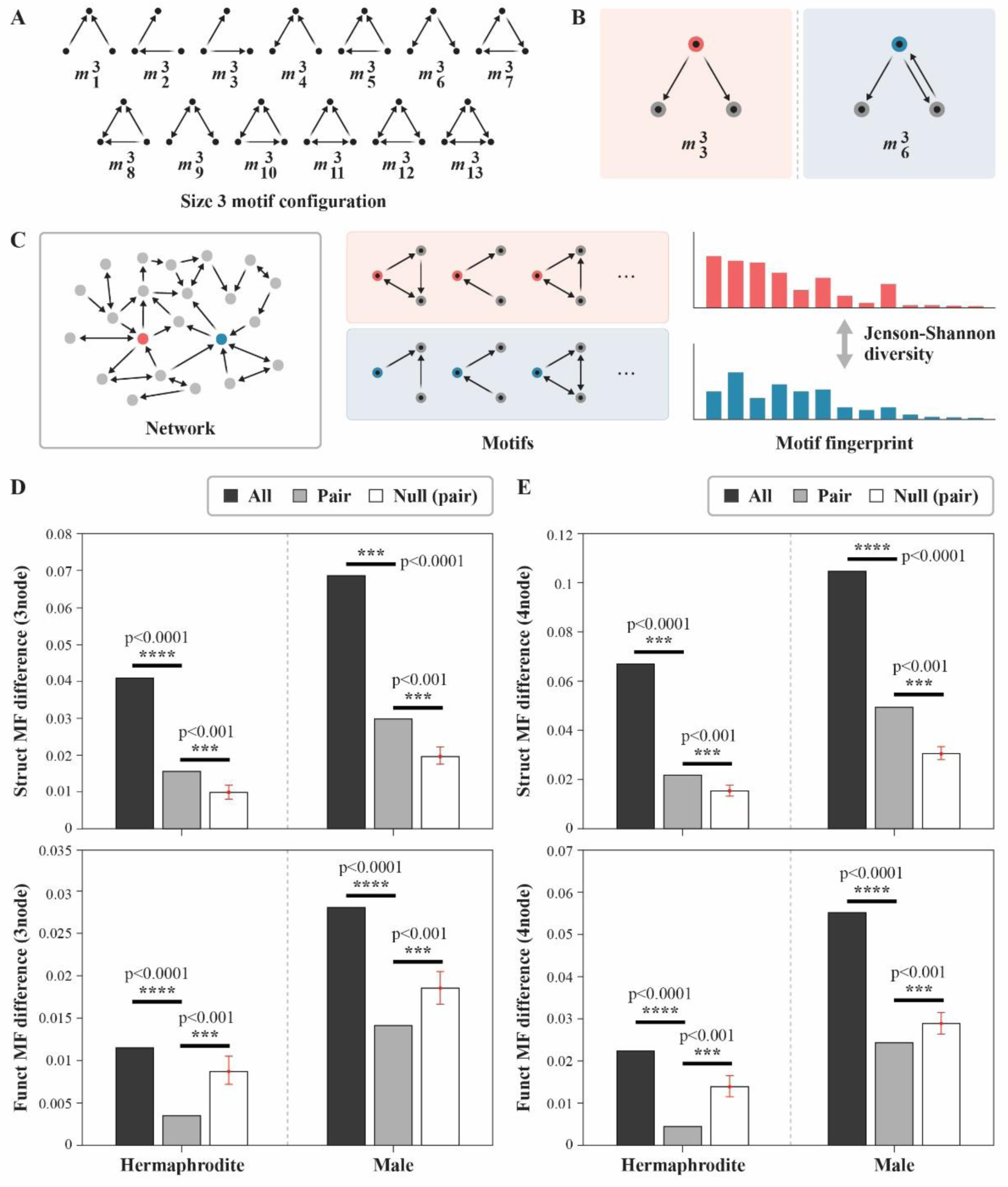
**(A)** All possible 3-node motif configurations. (B) **An example of a network that has a high mirror-symmetric ratio but functionally different motifs on each side.** Among those five edges, there is only one mirror-asymmetric edge on the right, resulting in a high mirror-symmetric ratio of 0.8. However, those two motifs, *m*^3^ and *m*^3^, have different circuitry properties: *m*^3^ exhibits synchronized activity but *m*^3^ does not. See main text for details. (C) **An illustration of how motif fingerprint (MF) difference is calculated.** The frequencies of motifs for each node are transformed into a motif probability distribution, and then Jenson-Shannon divergence is used to quantify the disparity of distributions (see Methods). (D) 3-node MF difference for *C. elegans* connectome and null models. (E) 4-node MF difference for *C. elegans* connectome and null models. In (D) and (E), the null-model result is the MF difference value between bilateral pairs in a randomly shuffled network, where the degree sequence is maintained.

In our motif analysis, we first computed the frequencies of 3-node and 4-node *structural motifs* for each node (Milo et al., 2002, Sporns and Kötter, 2004), then normalized these frequencies to transform them into a motif probability distribution (for further details, refer to the Methods section). To determine the difference in structural motif fingerprints between a pair of neurons, we utilized Jenson-Shannon (JS) diversity to compare the motif probability distributions of the given network (Figure 3C) (see Methods). This JS diversity value is what we refer to as the *motif-fingerprint difference* (MF difference). Our findings showed that the structural MF difference was significantly lower in bilateral neurons than in all neurons (p<0.0001 for all comparisons; see the upper panels in Figure 3D, E), indicating large bilateral redundancy in the network functions (as measured by motif structure). Intriguingly, when we compared those values with null models (i.e., random networks; see Methods), we found a significant discrepancy in the 3-node and 4-node structural motifs of bilateral neurons (indicated by a larger MF difference) compared to null models (see the upper panels in Figure 3D, E). In sum, although the structural motif fingerprints are more similar in the bilateral neurons than others, the asymmetric edges appear designed to make significant MF differences compared to null models.

To further understand these intricate findings, we conducted an additional analysis using a *functional motif*. A structural motif, which we have examined to this point, provides the anatomical structure for possible functional interactions between its comprising elements. However, it is important to note that in real-life neural networks, not all structural connections are always actively participating in functional interactions. This realization has introduced a different but related concept known as the *functional motif*, which targets function even more directly (Sporns and Kötter, 2004). If we consider structural motifs as the network’s physical infrastructure, functional motifs represent the variety of interactions that can occur within these structural boundaries, thus focusing more on the behavior of the elements. To elaborate, functional motifs are derived from a given structural motif and constitute the subset of motifs that can be formed by removing specific combinations of edges from the structural motif (see Methods). Using this concept, Sporns and Kötter (2004) have demonstrated that brain networks are configured to promote a high level of functional diversity (i.e., diverse functional motifs), leveraging a limited number of structural motif categories. This finding underscores the potential of functional motif to reflect the diverse functional roles a network can perform.

Building on previous work, we also calculated the functional motif fingerprint (MF) difference (i.e., MF difference using functional motif distribution; refer to Methods for more detail). First, as with the structural motifs, the functional MF differences were again significantly lower in bilateral neurons than in all neurons (p<0.0001 for all comparisons; see the lower panels in Figure 3D, E), indicating large bilateral redundancy in the network functions. Interestingly, however, we observed that the difference in functional motifs was significantly less than in our null models (lower panels in Figure 3D, E). This suggests that, even with substantial structural variations, the possible interactions exhibited by each pair of bilateral neurons (with each one’s neighbors) display remarkable similarity, indicating high bilateral redundancy. Given that the specific exclusion of asymmetric edges can lead to perfect symmetry in the motif configuration between bilateral neurons, this striking similarity in functional motif distribution is not surprising from a mathematical perspective. However, when paired with our earlier findings on structural motif differences, these results imply a crucial optimization between symmetry and asymmetry in structure: the significant structural differences can induce functional differences between bilateral neurons (i.e., functional lateralization), but the structures are also similar enough to enable similar functions. This, then, is a classic example of *degeneracy* — the ability of its diverse components to perform identical functions despite structural differences (Kitano, 2004, Tononi et al., 1999, Noppeney et al., 2004), which suggests a fine balance between functional diversity and biological robustness.

### Bilateral neurons exhibit high path-compensation

One potential consequence of having redundancy is the robustness of the nervous system (Dekker and Colbert, 2004, Tononi et al., 1999, Albert and Barabási, 2002). In particular, when two bilateral neurons (say A and B) are (at least partially) redundant regarding their functions, A can compensate B’s function (although not entirely as a backup) in case B is malfunctioning and vice versa (Kitano, 2004). In the previous studies on graph-theoretical vulnerability analysis of connectome, our team, along with other research groups, provided basic insight into the robustness (i.e., compensation capability) of the network (Gol’dshtein et al., 2004, Kim et al., 2016, Mishkovski et al., 2011). However, such vulnerability analyses generally focus on the effects of removing individual nodes or links from a network, measuring the impact on the system’s overall properties. While this approach provides valuable insight into the importance of individual components, it falls short of capturing the intricate interdependencies between pairs of neurons (Gol’dshtein et al., 2004, Kim et al., 2016, Mishkovski et al., 2011). Thus, we newly introduce a vulnerability-based measure, *path-compensation index*, to fill this analytical gap. This index quantifies the extent to which one neuron can compensate for another in the global information process when the latter is lost.

The path-compensation index for a specific neuron pair is defined as the discrepancy between the decline in global efficiency when both neurons are removed together and the linear summation of the decline in global efficiency when only one of them is removed individually (see Methods and Figure 4A). The global efficiency (which is the sum of inverse of shortest path lengths) is a measure that indicates how efficiently information is exchanged over the network (Rubinov and Sporns, 2010). If two neurons contribute independently to the global information process (for example, if they belong to two separate sub-networks), their simultaneous removal will cause a decrease in global efficiency equal to the sum of the decreases caused by removing each neuron individually. In this scenario, the path-compensation index would be zero, indicating no functional redundancy or dependency regarding the global information flow between these neurons. However, if there is a certain degree of redundancy between the roles of these neurons, i.e., one neuron can somewhat provide an alternate (i.e., redundant) pathway in the absence of the other, the combined decrease in global efficiency caused by removing both neurons individually would be less than their simultaneous removal. Thus, the path-compensation index is high when the impact of removing two neurons simultaneously is higher than the sum of impacts of removing each neuron individually – and so yielding a positive path-compensation index, implying functional redundancy or overlap. Conversely, if the removal of one neuron disrupts the function of the other (that is, eliminating one neuron disrupts certain pathways through the other), the effect of removing a single one can be close to when removing both. In this scenario, the summation of the individual decreases becomes more significant than the decrease observed when both neurons are removed, resulting in a negative path-compensation index. Hence, the path-compensation index quantitatively measures the functional redundancy and interdependence regarding the global information process between pairs of neurons in a network.

**Figure 4.**
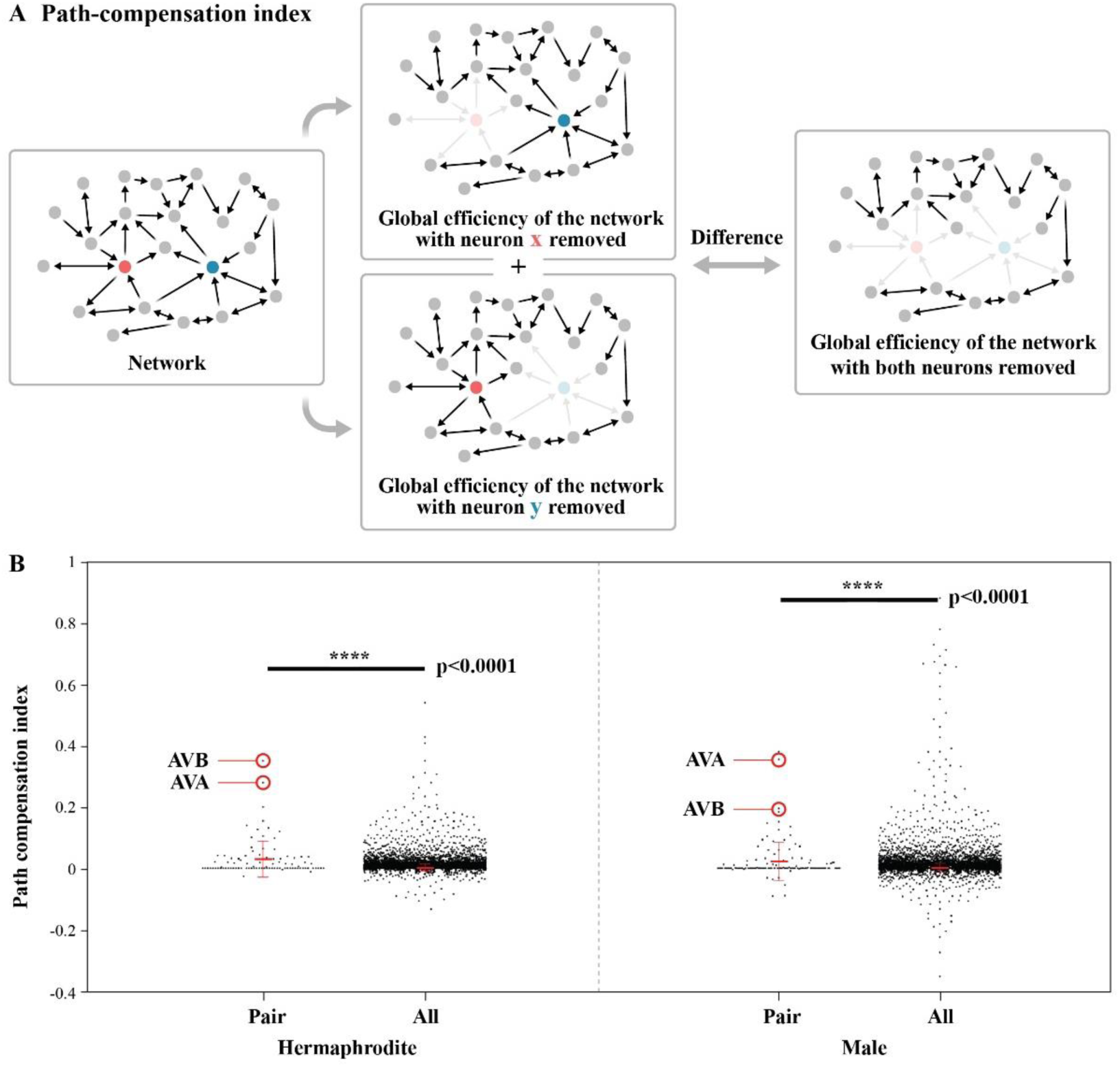
**(A)** An illustration of how path-compensation index is calculated. The path-compensation is defined as the difference between the decline in global efficiency when both neurons are removed together and the linear summation of the declines when they are removed individually. (B) **Path-compensation of *C. elegans* connectomes.** Bilateral pairs show a significantly higher path-compensation compared to all pairs in both sexes (*p* < 0.0001). The red error bar indicates the mean and standard deviation. The null-model result compares the path-compensation indexes between bilateral pairs in a randomly shuffled network, where the degree sequence is maintained, and bilateral pairs in the actual network of *C. elegans*. In both sexes, AVA and AVB neurons exhibit high path-compensation among bilateral pairs.

Based on this new measure, we found that the path-compensation index of bilateral neurons was significantly higher than all neurons (hermaphrodite: *p* < 0.0001, male: *p* < 0.0001; Figure 4). This finding implies that if one of the bilateral neurons is disrupted, its counterpart (i.e., contralateral neuron) can provide alternative pathways to maintain the system’s functionality, suggesting high redundancy between the bilateral neuron pair regarding the global information path. Notably, the interneuron pairs AVA and AVB were among the top 3 for pathway compensation in both hermaphrodite and male. These neurons are pivotal for controlling backward (reversal) and forward locomotion, respectively (as detailed in studies by Chalfie et al., 1985, Kawano et al., 2011, Wen et al., 2012). This result aligns well with previous findings that highlight the crucial role of AVA and AVB in driving robust initiating signals for movement (Kato et al., 2015, Kaplan et al., 2018, Uzel et al., 2022), which underscores the importance of robustness in the movement decision.

### Distinct level of redundancy in subsystems and individual neurons

Finally, we explored the *C. elegans* connectome in detail by investigating the redundancy within subsystems divided by three neuronal classes — sensory, motor, and interneurons (White et al., 1986). Our investigation involved the extraction of the first principal component (PC1) for each category of redundancy measures utilized in this study. Specifically, these measures encompassed connectivity similarity (including four specific measures: 1-step input, 1-step input-output, 2-step input, and 2-step input-output), MF difference (including four specific measures: 3-node structural, 3-node functional, 4-node structural, and 4-node functional), and path-compensation index (a single measure). We found that the PC1 accounted for the majority of the variance within each measure type, with values of 85% for hermaphrodite and 93% for male connectome in connectivity similarities, 83% for hermaphrodite and 90% for male in MF difference, and 100% for both sexes in path-compensation index (since there is only a single measure).

Given this significant observation, we used the PC1 values as representative indicators for each type of measure. Our analysis then progressed to a comparative evaluation of these PC1 values across the sensory, motor, and interneuron classes, allowing us to draw insights into the underlying redundancy patterns within these distinct subsystems of the *C. elegans* connectome. We found a consistent trend across the three types of redundancy measures (as depicted in Figure 5A-C). Sensory neurons typically displayed the lowest redundancy, characterized by the lowest connectivity similarity, the highest MF difference, and the lowest path-compensation index. In contrast, the redundancy of interneurons was the highest among the three classes in all the comparisons. Although the statistical results (i.e., p-values) were not consistent across the sexes and three categories of redundancy measures, significant differences were found in the hermaphrodite between the sensory and interneuron systems in all three measures (see the upper panels in Figure 5B, C). These differences remained significant even after correcting for multiple comparisons (p<0.05, Bonferroni correction, corrected for a total of three comparisons), indicating significant redundancy differences between sensory and interneuron classes. In addition, the motor-neuron system demonstrated a moderate level of redundancy, positioning itself between the sensory and interneuron systems (Figure 5A-C). Yet, it is important to acknowledge that our redundancy analysis was confined to bilateral neuron pairs, thus omitting unilateral neurons. Considering that the motor-neuron class in the *C. elegans* connectome consists of a substantial proportion of unilateral neurons (approximately 69% for both sexes, as seen in Figure 1A), the redundancy values derived from our analysis may not accurately represent the overall redundancy within the motor-neuron system. Therefore, instead of making an in-depth interpretation of such partial aspects of the motor system revealed in this analysis, it is noteworthy first to emphasize the unbiased nature displayed by the majority of unilateral neurons. A thorough discussion of this aspect, including potential interpretations, will be elaborated on in the Discussion section.

**Figure 5.**
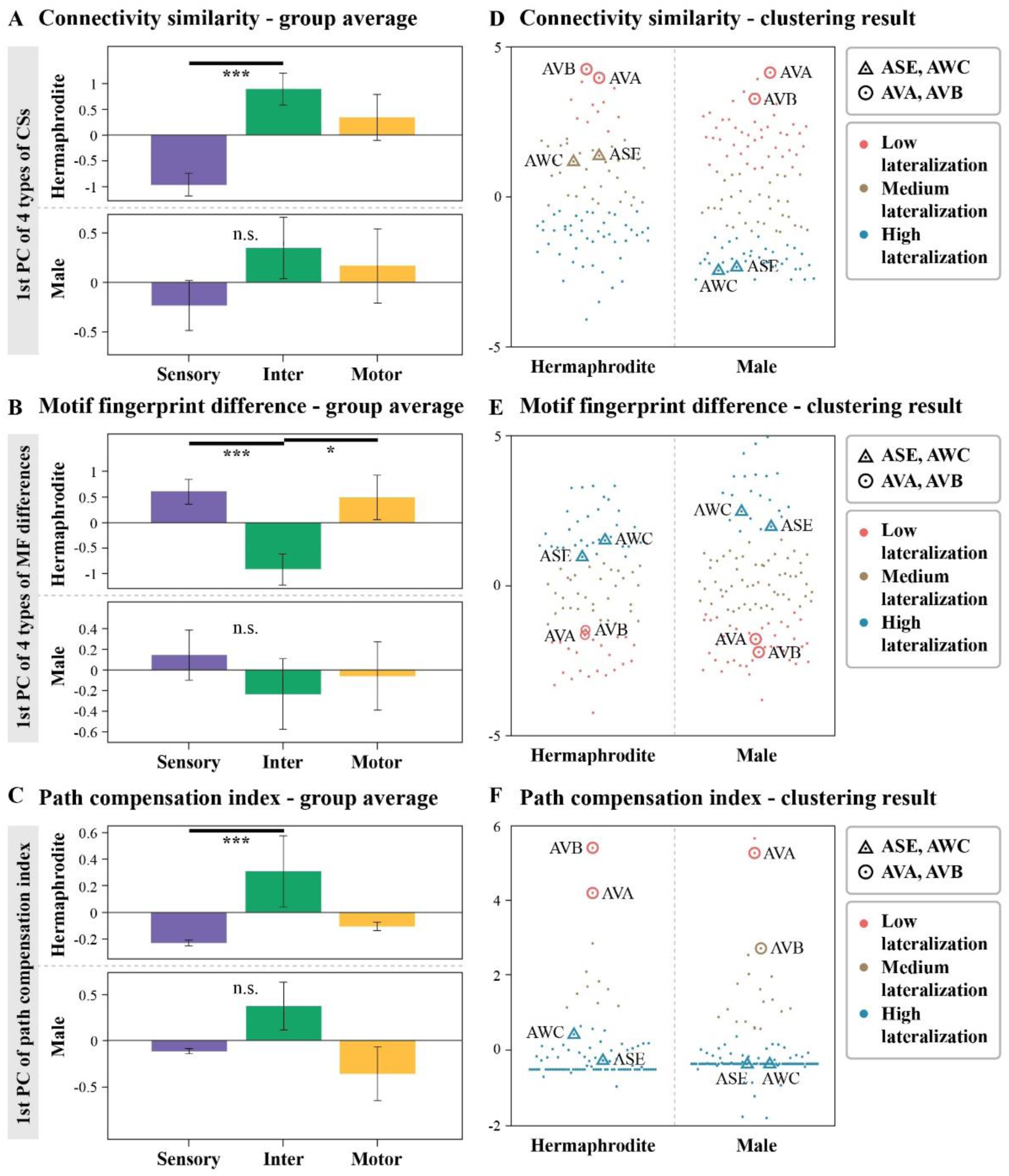
**(A)-(C)** Comparison of redundancy measures in the three subsystems: sensory, motor, and interneuron. In each category of redundancy measures, namely (A) connectivity similarity, (B) motif-fingerprint difference, and (C) path-compensation index, the first principal component (PC1) is extracted and utilized to evaluate and contrast redundancy among these subsystems. (D)-(F) **K-means clustering (K=3) analysis results.** This analysis categorizes the bilateral neurons into three distinct groups based on each category of redundancy measures: low lateralization (depicted in red), medium lateralization (in brown), and high lateralization (in blue). The diagrams also highlight well-known functionally lateralized neurons, such as ASE and AWC, as well as neurons AVA and AVB, which exhibit the highest path-compensation index.

Following this, we examined the redundancy of individual bilateral pairs, with a primary focus on the well-known functionally lateralized neurons, the ASE and AWC neurons, as well as the notable path-compensating neurons, the AVA and AVB neurons. We carried out three separate K-means clustering analyses (with K=3) using these distinct measures: one with four connectivity similarity differences, another with four MF differences (see Methods), and the other with path-compensation index. In each analysis, the bilateral neurons fell into three different groups, indicating low, medium, and high lateralization groups (Figure 5D-F). We then examined whether the functionally lateralized neurons, the ASE and AWC neurons, were all classified into the high lateralization group in both sexes. With the exception of one instance (connectivity similarity in hermaphrodites), the ASE and AWC neurons were consistently grouped into the highly lateralized group. This suggests that these functionally lateralized neurons indeed display a low degree of redundancy (i.e., a high degree of diversity). We also found that the significant path-compensating neurons, AVA and AVB (see Figure 4B), were all categorized into the low lateralization group in the analyses utilizing connectivity similarity or MF difference, indicating a substantial level of redundancy in AVA and AVB neurons’ functionalities as well as high level of path-compensation. The names of neurons categorized into low, medium, and high lateralization groups in each clustering analysis can be found in the Supplementary Material.

## Discussion

Investigating the bilateral symmetry and asymmetry of nervous systems promises to advance our understanding of its functionality and design objectives. In the current study, we attempted to unravel the degree of bilateral symmetry in the *C. elegans* connectome by introducing various graph-theoretic measures. The symmetry ratio reflected the great extent of structural symmetry in the *C. elegans* connectome of both sexes, and the various graph-theoretic measures (such as connectivity similarity, MF difference, and path-compensation index) all indicated bilateral redundancy in the functions of the left and right networks. Notably, a closer examination of the subsystems reveals distinct differences in the degree of symmetry between the sensory, motor, and interneuron systems from the perspective of bilateral redundancy and diversity. We next discuss our findings focusing on the following key aspects: (1) bilateral symmetry and redundancy in the *C. elegans* connectome, (2) bilateral degeneracy in the *C. elegans* connectome, and (3) design objectives of sensory, motor, and interneuron systems.

### Bilateral symmetry and redundancy in the C. elegans connectome

The *C. elegans* connectome exhibits a large degree of bilateral symmetry, evidenced by the high mirror-symmetry ratio, which might reflect significant redundancy within the nervous system, at least in certain aspects. Indeed, the redundancy measures highlight the extensive functional or operational overlap between the *C. elegans* connectome’s left and right sides. In this study, three redundancy measures were employed: connectivity similarity, MF difference, and path-compensation index. In each of the three measures, the bilateral neuron pairs consistently exhibit greater redundancy than any other possible neuron pairs within the network. It is important to note that all three measures can also be seen as alternative forms of measuring bilateral symmetry based on the different fundamental assumptions, each accommodating a specific level of abstraction that overlooks certain details. This multifaced approach provides a more detailed insight into symmetry and redundancy, which is particularly relevant given the distinct complexities and asymmetries found in the *C. elegans* connectome.

First, connectivity similarity focuses on the similarity in input information and output destinations without considering the overall network topology or specific local connectivity patterns. By abstracting away these details, connectivity similarity enables the identification of redundancies that pertain to the processing of specific information, providing a unique perspective on how neurons can be equivalent or similar to handling particular tasks or signals. This is supported by Uzel et al. (2022) in their structure-function predictions: specifically, the connectivity similarity was able to predict the neural dynamics. Therefore, the notable connectivity similarity among bilateral neuron pairs in the *C. elegans* connectome points to a potential symmetry in their functional processing.

Furthermore, motif-fingerprint difference takes a different approach by concentrating solely on local wiring patterns, disregarding global topology and the exact identity of input-output information. By focusing on these local patterns, MF difference can highlight similarities or differences in operations even when two neurons share the same signal inputs. Thus, our finding of the low MF difference in bilateral neuron pairs of the *C. elegans* connectome indicates that the possible operations between them are indeed similar.

Finally, path-compensation index captures the redundant role of neurons as channels for global information flow. It does not consider specific topological details or local connectivity, focusing instead on how global signals pass through the given pair of neurons. By ignoring these intricate details, this measure helps identify the overarching functional redundancy regarding the integration or relay of global information between neuron pairs. Our study has shown redundancy in this respect in the *C. elegans* connectome.

It is essential to recognize that the relation between bilaterally symmetric structure (i.e., bilateral connections) and redundancy is intricate. The optimization of redundancy should not be simplified as merely an increase in the ratio of mirror-symmetric edges since the asymmetric edges also contributed to enhancing bilateral redundancy (see Result section 3 as well as Supplementary figure 2). Instead, it represents a more intricate process that warrants further exploration in future studies.

### Bilateral asymmetry and degeneracy in the C. elegans connectome

Degeneracy is characterized as the capacity of various elements within a system to carry out the same function despite structural differences, which allows different parts of a system to show different functions but enables performing similar tasks if necessary (Kitano, 2004, Tononi et al., 1999, Noppeney et al., 2004). In the real biological context, where the duplication of identical elements for a single function can be costly, degeneracy may represent a feasible strategy to achieve robustness (Kitano, 2004). Put differently, if the bilateral neurons evolved via duplication, they then could potentially modify to accomplish different asymmetric function while maintaining some degree of redundancy (Pohl, 2011, Hobert, 2014). In our examination of the *C. elegans* connectome, we found that although the bilateral sides were highly redundant, none of the redundancy measures indicated the presence of entirely identical elements (e.g., with connectivity similarity equal to 1 or MF difference equal to 0) between them. Rather, we observed that approximately 33% of the connections were asymmetric, which exceeds the experimental limit of individual variance in connections (about 25%). (White et al., 1986, Varshney et al., 2011, Morone and Makse, 2019). This finding highlights significant diversity between the left and right sides of the *C. elegans* connectome. These observations provide partial support to the idea of bilateral degeneracy in the *C. elegans* connectome (Fornito et al., 2016).

The analysis based on MF difference adds more explicit support for this: when compared with null models, the structural motifs between bilateral sides were found to be significantly different, but the functional motifs were remarkably alike. This indicates that even with variations in structural aspects, the potential functions can be identical, revealing that degeneracy may indeed serve as a foundational design principle in this and likely other (at least bilateral) nervous systems. Moreover, the motif analysis in our study offers more profound insights into the intricate process of achieving degeneracy, a principle that may be fundamental in neural design. Given that there are only minor proportions of asymmetric edges, identical local processing could be efficiently realized by emulating the same local network topology with only minor alterations to asymmetric connections. Such inactivation could be temporary, enhancing robustness against temporary disturbances or malfunctions, or long-standing for a rewiring strategy to compensate for permanent unilateral damage. Numerous previous connectome studies highlight that a core design principle of the nervous system is to minimize wiring costs, i.e., the energy consumption involved in forming or maintaining connections (Cherniak, 1994, Chklovskii et al., 2002, Klyachko and Stevens, 2003, Rivera-Alba et al., 2011). Thus, the possibility of emulating contralateral local topologies with minimal rewiring aligns well with the goal of efficient energy use.

Such a principle, if applicable to other species, offers an appealing explanation for the frequently observed contralesional compensation in patients with unilateral hemispheric damage: for example, one patient with left cerebral infarction recovered his sensory and motor functions, attributed to an enhanced connection between the right side of his brain and his right limbs (Shi and Wan, 2021); a study with 15 participants revealed significant plastic changes, an indication of altered motor-output organizations, in the unaffected hemisphere post-ischemic stroke (Netz et al., 1997); and other studies have reported right-hemispheric compensation for language function in patients recovering from stroke-induced aphasia (Crinion and Price, 2005, Mimura et al., 1998, Jodzio et al., 2005, Karbe et al., 1998). From these observations, it is possible to infer that the contralesional compensation seen in cases of unilateral brain damage may be a direct consequence of the brain’s remarkable bilateral symmetry, working in tandem with the principle of minimizing wiring costs. This combination allows for a flexible response to injury, promoting recovery and adaptation by leveraging existing structural principles within the nervous system.

### Design objectives of sensory, motor, and interneuron systems

Our study reveals distinct redundancy profiles of subsystems of the *C. elegans* connectome, characterized by prominent asymmetry in the sensory system (indicating lateralization) and prominent symmetry in the interneuron system (indicating redundancy). While the statistical significance was not consistent across sexes and specific measures (as depicted in Figure 5A-C), we speculate that the redundancy profile (i.e., high lateralization in sensory neurons and high redundancy in interneurons) could be indicative of the unique functional objectives intrinsic to each system since the redundancy trend was generally consistent and the difference between sensory neurons and interneurons remained statistically significant after multiple-comparison correction (refer to Results for details).

First, the asymmetric property of sensory neurons indicates a strategy to diversify the information type the individuals receive from the environment. Note that the well-known functional lateralized neurons in the *C. elegans* connectome (i.e., ASE and AWC) are all sensory neurons. By diversifying the types of information an individual can gather from their surroundings, the organism may better detect and exploit available resources (Jordan and Ryan, 2015), potentially improving its survival and enabling more efficient location of mates (Arnqvist, 2006, Ryan, 1990) or detection of predators (Jordan and Ryan, 2015). Therefore, organisms that can perceive a wider spectrum of environmental signals often possess an evolutionary advantage in terms of overall adaptability, which is in-line with the importance of having asymmetry in the sensory system.

Conversely, interneurons in *C. elegans*are crucial for directing a variety of behavioral patterns and extensively interacting with other neurons (Hendricks et al., 2012, Ghosh et al., 2017, Gordus et al., 2015), including sensory and motor neurons, necessitating a robust operational framework for signal transmission and integration. The high symmetry in the interneuron system enables such robust operation since redundancy results in increased resilience to internal signal noise or random perturbation and even robustness to partial damage of networks (Dekker and Colbert, 2004, Tononi et al., 1999, Albert and Barabási, 2002). Also, the current study has shown that two interneurons, the AVA and AVB neurons, exhibit high redundancy, well in line with the previous discussion that those neurons are crucial to robust locomotion-related decision (Kato et al., 2015, Kaplan et al., 2018, Uzel et al., 2022). Moreover, the high symmetry of interneuron connections can also serve an essential role in integrating the diverse and asymmetric information gathered by sensory neurons. The bilateral symmetry in the interneuron system allows for robust and balanced processing of these diverse inputs from asymmetric sensory neurons, then turning them into meaningful signals for unbiased behavioral responses and keeping a balance between both sides of the organism.

While our analysis reveals that the motor-neuron system maintains a moderate level of symmetry between sensory and interneuron systems, we should note that this investigation primarily focused on bilateral neurons. In other words, this approach does not consider a significant portion of the motor-neuron system, since most of these motor neurons are unilateral neurons (about 69% for both sexes), positioned on the medial and ventral parts of the organism’s body (see Figure 1A). Given this predominance of unilateral and medially aligned neurons, the motor-neuron system exhibits a highly unbiased structure, possibly resulting in unbiased functionalities. Moreover, by considering unilateral neurons as either merged pairs at the center or as neurons that have not evolved into symmetric pairs, we can adjust our methodology to apply the redundancy measures to these neurons, which could lead to a perfect redundancy and symmetric score. This implicates that the motor neuron system of *C. elegans* might exhibit highly symmetric and unbiased functionalities, which is one of the possible solutions for enabling unskewed directed locomotion (see Ruppert et al., 2004 for the evolutionary advantages of unskewed directed locomotion).

Nonetheless, the observed moderate bilateral asymmetry in some of the motor neuron pairs of *C. elegans* suggests a degree of complexity in its motor functions. This could potentially explain the asymmetrical behaviors seen in the mating rituals of male *C. elegans* (Downes et al., 2012). Although the results regarding the motor system were not statistically significant, making definitive conclusions challenging, we propose that future research should investigate this aspect further. Such studies would contribute to a more thorough comprehension of the behavioral asymmetries commonly found in animals (e.g., handedness).

### The role of bilateral symmetry and asymmetry in the sensory integration for coordinate motor outputs

This study provides crucial insights into the neural circuitry of *C. elegans*, emphasizing a strategic design principle from sensory inputs to motor outputs. In the *C. elegans* network, the neural circuit shows a consistent pattern. Sensory neurons (S) connect to 1-3 layers of interneurons (I), with the final layer of interneurons known as command interneurons. These command interneurons directly control motor neurons, which regulate muscle activity. Overall, this displays an S → I → M pattern (Gray et al., 2005, Jarrell et al., 2012, Qian et al., 2011). Regarding the bilateral symmetry, such a pattern can be interpreted as the following: “Asymmetric (diverse inputs)” → “Symmetric (robust integration)” → “Unbiased (balanced coordination).” Thus, this symmetry-asymmetry interplay between subsystems could be key to enabling complex and adaptive behaviors, highlighting the sophistication of neural design strategies in *C. elegans*.

Our exploration leverages the directed graph model to dissect the topological nuances of bilateral redundancy, acknowledging the profound impact of network topology on neural dynamics (Uzel et al., 2022, Bullmore and Sporns, 2009). However, we recognize the model’s simplification of multifaceted biological interactions: The reality of multiple synapses between neuron pairs, featuring varying strengths and both excitatory and inhibitory interactions, suggests an additional complexity layer not captured in our current analysis. Future research should delve into these synaptic details to enrich our understanding of functional redundancy and the connectome’s robustness. Moreover, our study’s static analysis of the connectome underscores the need to consider the dynamic interplay within neuronal networks, where the patterns of interactions and signal propagation evolve. Incorporating dynamic analyses could unveil how structural characteristics identified in our study manifest in the network’s functional activities over time, offering a more nuanced view of the connectome’s operational principles (Tononi et al., 1999, Uzel et al., 2022, Chapman and Mesbahi, 2015, Whalen et al., 2015, Yuan et al., 2013, Randi et al., 2023). Although integrating dynamic elements poses challenges due to increased complexity, it represents a promising avenue for deepening our comprehension of neural network functionality.

In navigating from topological insights to a comprehensive grasp of dynamic neural interactions, our study lays the groundwork for subsequent explorations. It highlights the critical role of bilateral symmetry and asymmetry within neural architectures in fostering adaptability in behavior. This insight, while rooted in the study of *C. elegans*, may extend its relevance to wider neuroscientific and computational studies, suggesting universal principles of neural design.

## Methods

### Constructing directional graphs for male and hermaphrodites C. elegans

Our study involves an analysis of the most recent connectome data pertaining to adult hermaphrodites and male *C. elegans,* as presented in the work conducted by Cook and his colleagues (Cook et al., 2019). The detailed depiction of connections within this connectome data encompasses various attributes, including the nature of connections (chemical synapses or gap junctions), their direction, and their strength, as quantified by the number and sizes of connections. For the sake of this paper, we excluded the pharyngeal neurons from the connectome data for both genders due to their distinction from the somatic nervous system.

We combined the network of gap junctions and chemical synapses to generate directed and binary networks for *C. elegans*. Despite the differences between gap junctions and chemical synapses, our emphasis lies in concentrating on the process of information transfer, with a deliberate disregard for the specific characteristics and intensities of these connections. As a result, we were able to establish connectomes comprising 4,916 connections among 280 neurons in hermaphrodites and 5,490 connections among 362 neurons in male. Both genders share 272 neurons, alongside sex-specific neurons unique to each gender (8 in hermaphrodite and 90 in male). Additional details concerning individual *C. elegans* neurons, including their functions, were extracted from the wormatlas website (Altun and Hall, 2015).

The *C. elegans* connectomes are graphically represented as *G* = (*V*, *E*), where *V* represents the set of nodes (neurons *x*, *y*, …) and *E* represents the set of edges (connections (*x*, *y*),)) among these nodes. The connectivity structure is succinctly summarized by the adjacency matrix A, wherein *A*_*ij*_ = 1 if (*i*, *j*) ∈ *E* which signifies a direct connection from neuron *i* to neuron *j*, while *A*_*ij*_ = 0 if (*i*, *j*) ∉ *E* which denotes the absence of such a connection. Self-connections are treated as nonexistent (∀*A*_*ii*_ = 0).

### Mirror-symmetry index

The left-right symmetry of *C. elegans* neurons has been comprehensively categorized in cellular contexts through the efforts of White and his colleagues. By utilizing electro-macroscopy data to reconstruct the nervous system of *C. elegans*, White grouped neurons with analogous structures into distinct classes, denoting their symmetry through the final letter of the neuron’s name as “L” or “R” (for instance, the AVA neuron class consisting of AVAL and AVAR). Roughly two-thirds of *C. elegans* neurons (198 out of 302 neurons in hermaphrodites and 250 out of 385 neurons in males) exhibit bilateral symmetry. The remaining neurons are predominantly situated on or near the midline and lack contralateral counterparts.

Derived from these anatomical symmetry attributes in *C. elegans*, we can define contralaterality, marked as (*), as the bilateral counterpart of a given component. For instance, within a bilaterally symmetric neuron pair, if a neuron resides on the left side of the body, its contralateral partner in the pair would be positioned on the right side (for example, *x*=ASEL is the contralateral neuron for *x*\*=ASER). Unlateral neurons lack corresponding contralateral counterparts, thus resulting in defining contralateral neurons of unlateral neurons as the neurons themselves (for instance, the *u*=CA01 neuron serves as its own contralateral neuron, *u*\*=*u*=CA01). The existence of contralateral neurons inherently introduces the concept of establishing a contralateral edge or link between them. Specifically, we introduce the term “contralateral edge,” symbolized as (*), to represent the connection between corresponding contralateral neurons.

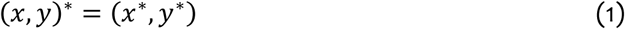

A mirror asymmetric edge is identified by its presence when the contralateral edge, which connects corresponding contralateral neurons, is absent within the network. Conversely, a mirror symmetric edge is defined as a link that manifests when a contralateral edge exists between the corresponding contralateral neurons.

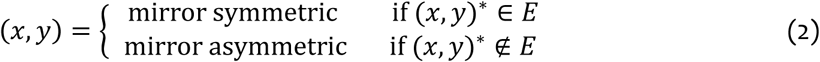

### Connectivity similarity

The connectivity similarity measure is designed to determine the degree of resemblance between the sets of neighboring nodes, facilitating the detection of potential symmetry in the connectivity identities. In this study, we focused on the two types of distinction of neighborhoods: the 1-step/2-step neighborhood and the input/output neighborhood. The 1-step/2-step neighborhood includes nodes that are connected to the given node by either one or two directed steps. Meanwhile, the input and output neighborhood consist of nodes that provide inputs to the node under consideration and those that receive outputs from it, respectively. The neighborhood of neuron *N*(*x*) is formally defined as follows:

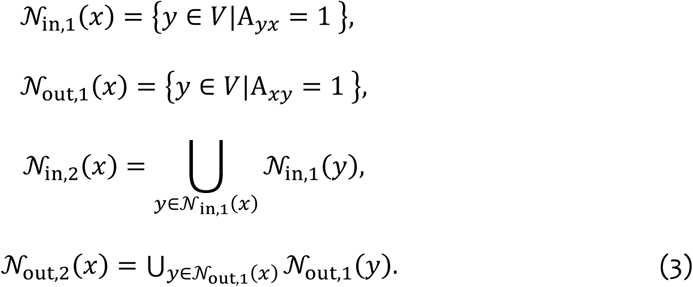

With the definition of neighborhood, *k*-step input connectivity similarity (CS_in,*k*_) and *k*-step input-output connectivity similarity (CS_in−out,*k*_) is defined as follows:

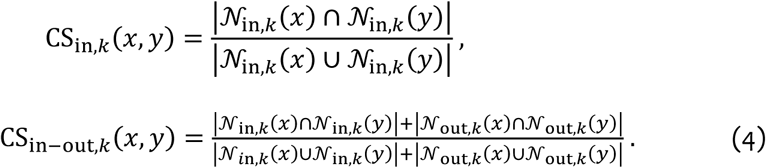

### Motif-fingerprint (MF) difference

Motifs are defined as local connectivity patterns comprising a set of *n* nodes, without any fragmented components or nodes, and they form the core elements of a network’s architecture (Milo et al., 2002, Sporns and Kötter, 2004). For instance, in the case of triads (motif size 3), there exist 13 potential configurations (see Figure 3), whereas for tetrads (motif size 4), the number of possible configurations rises to 199. Our investigation analyzed normalized motif fingerprints, mainly focusing on 3 and 4-node structural and functional motifs. The motifs fall into two main categories: structural and functional. Structural motifs are the local patterns of physical connections. In contrast, functional motifs include all possible non-fragmented subgraphs of these local connectivity patterns found within structural motifs, thus representing the possible realization of elementary operation using given structural motifs.

The MF difference was developed to measure the disparity between the motif probability distributions (be they structural or functional) in which a particular pair of neurons is involved. To determine the *n*-node motif probability distributions for a specific neuron *x* (denoted as a probability vector *p*→^*p*^_x_, with each vector component indicating the probability of the *k*-th motif configuration around neuron *x*), we first calculated the *n*-node motif frequency around neuron *x* (*m*→→→^*p*^, where each vector component corresponds to the frequency of the *k*-th motif configuration). This calculation was performed using the Brain Connectivity Toolbox (Rubinov and Sporns, 2010). We then transformed this motif frequency into a probability distribution, applying Laplace (add-one) smoothing to avoid the problem of zero probability. The formula for the *n*-node motif probability distributions for neuron *x* is as follows:

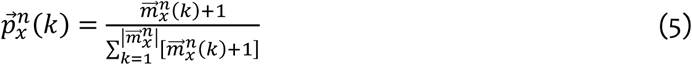

where *m*→→→^*p*^(*k*) and *p*→^*p*^(*k*) are the *k*-th components of *m*→→→^*p*^ and *p*→^*p*^ (i.e., corresponding to the *k*-th motif configuration), respectively. Then, the MF difference between neuron *x* and neuron *y* was defined as the Jenson-Shannon divergence (*D*_JS_) between the motif probability distributions of the given neurons *x* and *y*: *D*_JS_(*p*→^*p*^ ∥ *p*→^*p*^). This same method was used to compute the four variants of MF difference, specifically the 3-node structural, 3-node functional, 4-node structural, and 4-node functional differences.

### Path-compensation index

We define path-compensation index to quantify the extent to which one neuron can substitute for another in the overall process of information transmission, in cases where the latter is compromised (essentially, providing a redundant pathway). To calculate the path-compensation index for neurons x and y, we initially calculate the vulnerability of removing a set of neurons *Y* concerning all paths between any two neurons in *X*. The formula for vulnerability is given as follows:

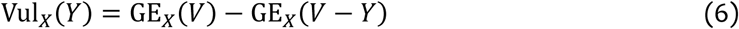

where GE_*X*_(*Z*) is the sum of the inverses of the shortest paths between any two neurons in *X* when only edges connected between the neurons in *Z* are available for traveling paths. Thus, Vul_*X*_(*Y*) represents the vulnerability of the paths (i.e., the extent of alteration required in the shortest paths) between any two neurons in *X* when the neurons in *Y* (as well as the edges connected with the neurons in *Y*) are removed from the graph G. Then, path-compensation index (PCI) for neuron *x* and *y* is defined as the discrepancy between the vulnerability of the paths between any neurons in *V* − {*x*, *y*} when both neurons are removed together (i.e., Vul_*V*−{*x*,*y*}_({*x*, *y*})) and the linear summation of the vulnerabilities of the paths between any neurons in *V* − {*x*, *y*} when only one of them is removed individually (i.e., Vul_*V*−{*x*,*y*}_({*x*}) + Vul_*V*−{*x*,*y*}_({*y*})). Thus, PCI for neuron *x* and *y* is defined as follows:

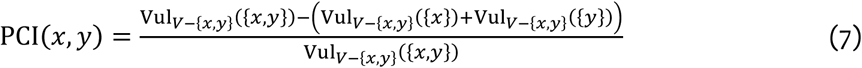

where the value is normalized by Vul_*V*−{*x*,*y*}_({*x*, *y*}) to ensure that the PCI becomes one when the input-output connectivity similarity between neuron *x* and *y* is one (i.e., when neuron *x* and *y* connected with exactly identical neurons).

### Null models for connectome (i.e., random networks)

In the current study, we employed the Maslov-Sneppen rewiring algorithm to generate null models that uphold the original network’s degree distribution. Preserving this degree distribution was pivotal in maintaining the inherent structural attributes of the network, even as we perturbed its fundamental connectivity arrangements. Subsequently, we subjected the network to approximately 1000 rewiring iterations for each connection. In total, we created 1000 null model networks to ensure robust statistical validation.

### Symmetry breaking and asymmetry breaking networks

Additionally, we utilized an alternative null model to examine the impact of both symmetric and asymmetric edges on redundancy measures within the network. The primary objective of this model was to determine whether network metrics predominantly derive from symmetric or asymmetric edges. To specifically investigate the influence of mirror symmetry, we adjusted the proportions of asymmetric edges using the Maslov-Sneppen rewiring algorithm while ensuring that the remaining asymmetric edges and all symmetric edges were preserved. Conversely, we conducted a complementary procedure by manipulating the proportions of symmetric edges while retaining the remaining symmetric edges and all asymmetric edges. Notably, during the shuffling of symmetric edges, contralateral edges were rearranged concurrently.

This iterative process was carried out across a range of proportions, varying from 0 to 100 percent in increments of 10 percent. For each proportion, the corresponding fraction of edges (either symmetric or asymmetric) was randomly chosen and shuffled with maintenance of degree distribution, then combined with the remaining unselected (i.e., unshuffled) portion of the network. For instance, in the case of 30 percent mirror symmetric shuffling, we randomly chose 30 percent of the symmetric edges and shuffled them using the Maslov-Sneppen rewiring algorithm. This created a new network comprised of the selected symmetric edges (the 30 percent). After this, the unselected symmetric edges and asymmetric edges were preserved and recombined with the shuffled selected network. This process was repeated to generate 100 randomized networks for each specified proportion.

### Custom shuffle test for comparison between bilateral and all possible neuron pairs

When performing statistical tests between bilateral pairs and all possible neuron pairs, the network measure values might not be statistically independent, as required by certain conventional statistical tests. To address this, we used a specialized resampling technique that does not depend on specific assumptions about the underlying data distribution. In this custom shuffle test, we randomly selected neuron pairs in number equal to the original bilateral neuron pairs and computed the Welch’s t statistics between this sampled pair group and all neuron pairs. This process was repeated 100,000 times. For the p-value estimation, we calculated the proportion of these absolute t-values that were equal to or greater than the original t-value’s absolute value, which was determined between the original bilateral neuron pairs and all neuron pairs.

### Subsystem analysis

For analyzing the difference of redundancy between subsystems (i.e., sensory, motor, and interneurons), we extracted the first principal component (PC1) for each category of redundancy measures (connectivity similarity – four measures, MF difference – four measures, and path-compensation index – one measure; detailed in the main text). Note that for the MF difference measure, we applied a logarithmic transformation prior to principal component analysis (PCA) to correct the skewed distribution and restore normality in the data. In addition, before performing PCA, the graph-theoretic measures were standardized, ensuring each measure had a mean of zero and a standard deviation of one, which guaranteed uniform variance across all variables. The choice to use the first principal component (PC1) was informed by its ability to represent the largest portion of variance in the dataset (detailed in the main text).

Employing the first principal component (PC1), we executed an additional custom shuffle test (for each category of redundancy measures). Initially, we computed Welch’s t statistics between every two groups. Subsequently, we randomly shuffled the group identity of each bilateral neuron pair (determining whether a pair is part of the sensory, motor, or interneuron systems). We ensured that bilateral neurons consistently grouped in identical subsystem. In each iteration of this process (each shuffle), we recalculated Welch’s t statistics for any two groups and collected the highest absolute t-value obtained. This procedure was repeated across 100,000 trials. To estimate the p-value, we determined the proportion of these maximum absolute t-values that were equal to or greater than the original t-value calculated between the group.

### Clustering bilateral neuron pairs

To distinguish each of bilateral neuron pairs into high, medium, low lateralization group, we employed K-means (K=3) clustering analysis for each category of redundancy measures (connectivity similarity – four measures, MF difference – four measures, and path-compensation index – one measure). Again, for the MF difference measure, we applied a logarithmic transformation prior to the clustering analysis. Also, before conducting the clustering analysis, the graph-theoretic measures were standardized, ensuring each measure had a mean of zero and a standard deviation of one, which guaranteed uniform scaling across all variables.

## Contributions

P.K. and J.J. initially conceived the study. P.K., Y.S., and J.J. designed and planned the study. P.K. and Y.S. developed analysis measures and produced the results. P.K. and Y.S. prepared the first draft, and all authors reviewed the manuscript. J.J. supervised the study.

## Ethics declarations Competing interests

The authors declare no competing interests.

